# Microtubules self-repair in living cells

**DOI:** 10.1101/2022.03.31.486545

**Authors:** Morgan Gazzola, Alexandre Schaeffer, Benoit Vianay, Jérémie Gaillard, Laurent Blanchoin, Manuel Théry

## Abstract

Microtubule self-repair has been studied both in vitro and in vivo as an underlying mechanism of microtubule stability. The turnover of tubulin dimers along the microtubule network has challenged the pre-existing dogma that only growing ends are dynamic. However, although there is clear evidence of tubulin incorporation into the shaft of polymerized microtubules in vitro, the possibility of such events taking place in living cells remains uncertain. In this study, we investigated this possibility by microinjecting purified tubulin dimers labeled with a red fluorophore into the cytoplasm of cells expressing GFP-tubulin. We observed the appearance of red dots along pre-existing green microtubule network within minutes. We found that the fluorescence intensities of these red dots were inversely correlated with the green signal, suggesting that the red dimers were incorporated into the microtubules and replaced the pre-existing green dimers. We then characterized the size and spatial frequency of these incorporations as a function of injected tubulin concentration and post-injection delay. The saturation of these measurements contradicted the hypothesis of nonspecific adsorption along microtubules and suggested that the injected dimers incorporated into a finite number of damaged sites. By our low estimate, within a few minutes of the injections, free dimers incorporated into major repair sites every 70 micrometers of microtubules. Finally, we mapped the location of these sites in micropatterned cells and found that they were more concentrated in regions where the actin filament network was less dense and where microtubules exhibited greater lateral fluctuations. These results provide evidences that microtubules do self-repair in living cells, and they provide a quantitative characterization of the temporal and spatial dynamics of this process in PtK2 cells.

## Introduction

Microtubules are polar and dynamic polymers running through the cytoplasm of eukaryotic cells. They serve as tracks for molecular motors, and as such play essential roles in key cellular processes such as cell polarity, migration, division and more generally to the global regulation of intracellular organization (Goodson & Jonasson, 2018). They result from the self-assembly of tubulin dimers into twelve to fourteen protofilaments that align longitudinally to form a hollow tube. Tubulin dimers are thus densely packed and highly organized into a pseudo crystallin structure named the microtubule lattice (Goodson & Jonasson, 2018). Microtubules dynamics is characterized by the alternance between growing and shrinking phases resulting from the addition and removal of tubulin dimers at the end of microtubules (Mitchison & Kirschner, 1984; Brouhard & Rice, 2018). However, in the last decade, it has been proposed that tubulin dimers could also be exchanged along the microtubule shaft in a self-renewal process of the lattice (Cross, 2019; Théry & Blanchoin, 2021).

Microtubules that were polymerized *in vitro* and further capped to prevent ends shortening were found to form kinks following the removal of free surrounding tubulin (Dye *et al*, 1992). Interestingly, these microtubules were capable to recover their linear shape after the re-addition of free dimers (Dye *et al*, 1992). This was suggestive of the loss and reincorporation of tubulin dimers in the lattice of microtubules, an hypothesis further demonstrated by the visualization of this process with fluorescent tubulin dimers (Schaedel *et al*, 2019). These exchanges were suggested to occur at specific defect sites where the lattice structure displayed defects such as missing dimers or dislocation in the organization of protofilaments (Chrétien & Fuller, 2000; Chretien *et al*, 1992; Guyomar *et al*, 2021; Schaedel *et al*, 2019). This hypothesis is consistent with the analysis of microtubule response to bending cycles during which they appeared to soften in a non-elastic response, and were further seen to recover their initial mechanical state by the incorporation of new dimers in their damaged lattice (Schaedel *et al*, 2015).

Tubulin dimers were even seen to incorporate into large sections in the shaft of taxol-stabilized microtubules (Reid *et al*, 2017). Such large incorporations of free dimers in damaged lattice have also been observed in response to severing enzymes activity (Vemu *et al*, 2018; Zanic, 2021), oxidative stress (Goldblum *et al*, 2020), and even the walk of molecular motors (Triclin *et al*, 2021). Synergistically, specific microtubule-associated proteins could either promote the incorporations of tubulin dimers, or act as a scaffold to protect damaged microtubule lattice (Aher *et al*, 2020; Lawrence *et al*, 2021). Overall, last *in vitro* studies challenged the rules of microtubule dynamics being restricted to the end, and redefined the lattice as a dynamic and self-repairing structure.

Unfortunately, in living cells, the investigation of microtubule self-repair has been more challenging. Experiments with photoconvertible tubulin were limited by the weakness of the signal, and the formation of aggregates that could incorporate during microtubules polymerization (Aumeier *et al*, 2016). Moreover, even though structural defect or larger damaged regions were undoubtedly observed in the microtubule lattice of mammalian cells by cryo-electron microscopy (Atherton *et al*, 2018; Chakraborty *et al*, 2020; Atherton *et al*, 2021), the incorporation of tubulin dimers in these regions has never been documented. Physical injuries or molecular motors overexpression were recently shown to increase microtubule length and network density in migratory cells and thus were suggestive of a self-repair mechanism (Andreu-Carbó *et al*, 2021; Aumeier *et al*, 2016). In addition, the entry of small peptides in the microtubule’s lumen implied the presence of sufficiently large opening in the microtubule lattice (Nihongaki *et al*, 2021). Finally, markers of tubulin dimers in a GTP-like conformation, i.e., newly added tubulin or locally damaged lattice, such as hMB11 or CLIP-170 were observed along the length of microtubules living cells (Dimitrov *et al*, 2008; de Forges *et al*, 2016). Although these results were suggestive of microtubule self-repair in living cells, they did not constitute clear evidence of its actual occurrence, nor they provide any quantitative description of its manifestation.

Here, to investigate the potential incorporation of tubulin dimers in pre-existing microtubules, we microinjected fluorescently labeled tubulin dimers in the cytoplasm of living cells. Our data demonstrate that injected dimers were mostly recruited to microtubule growing ends, but also within the microtubule lattice in a specific manner.

## Results

### Free tubulin dimers can form patches along pre-existing microtubules

Microinjection of biotin-bound tubulin dimers was used in early studies to investigate the localization of microtubules polymerizing sites (Schulze & Kirschner, 1986, 1987). Immuno-labelling of endogenous and injected dimers clearly revealed the addition of dimers at the end of microtubules, but this technique may not provide the required specificity to undoubtedly characterize repair sites. To circumvent the use of immunolabeling, we microinjected purified tubulin labelled with ATTO-565, to which we further referred to as injected tubulin (**IT**) (displayed in figures in magenta) in the cytoplasm of PtK2 cells expressing GFP-tubulin, to which we further referred to as expressed tubulin (**ET**) (displayed in figures in green) (**Figure 1a**). As expected from previous studies (Schulze & Kirschner, 1986), about half of the microtubule network got renewed in the 4 minutes following microinjection of 10µM of tubulin dimers (**Figure 1b**). The visualization and evaluation of potential repair sites appeared challenging in such a densely labelled network. We thus analyzed the tubulin content of pre-existing microtubules only 2 minutes after the microinjection (**Figure 1c**). Interestingly, we could observe the presence of patches of IT in several microtubules despite the short delay after the injection (**Figure 1d**, white arrows). However, the presence of those patches distant from growing ends, was insufficient to ascertain their origin. Indeed, they could result from IT incorporation within the lattice, IT binding along the side of microtubule shaft or even the annealing between two microtubules growing ends.

**Figure 1:**
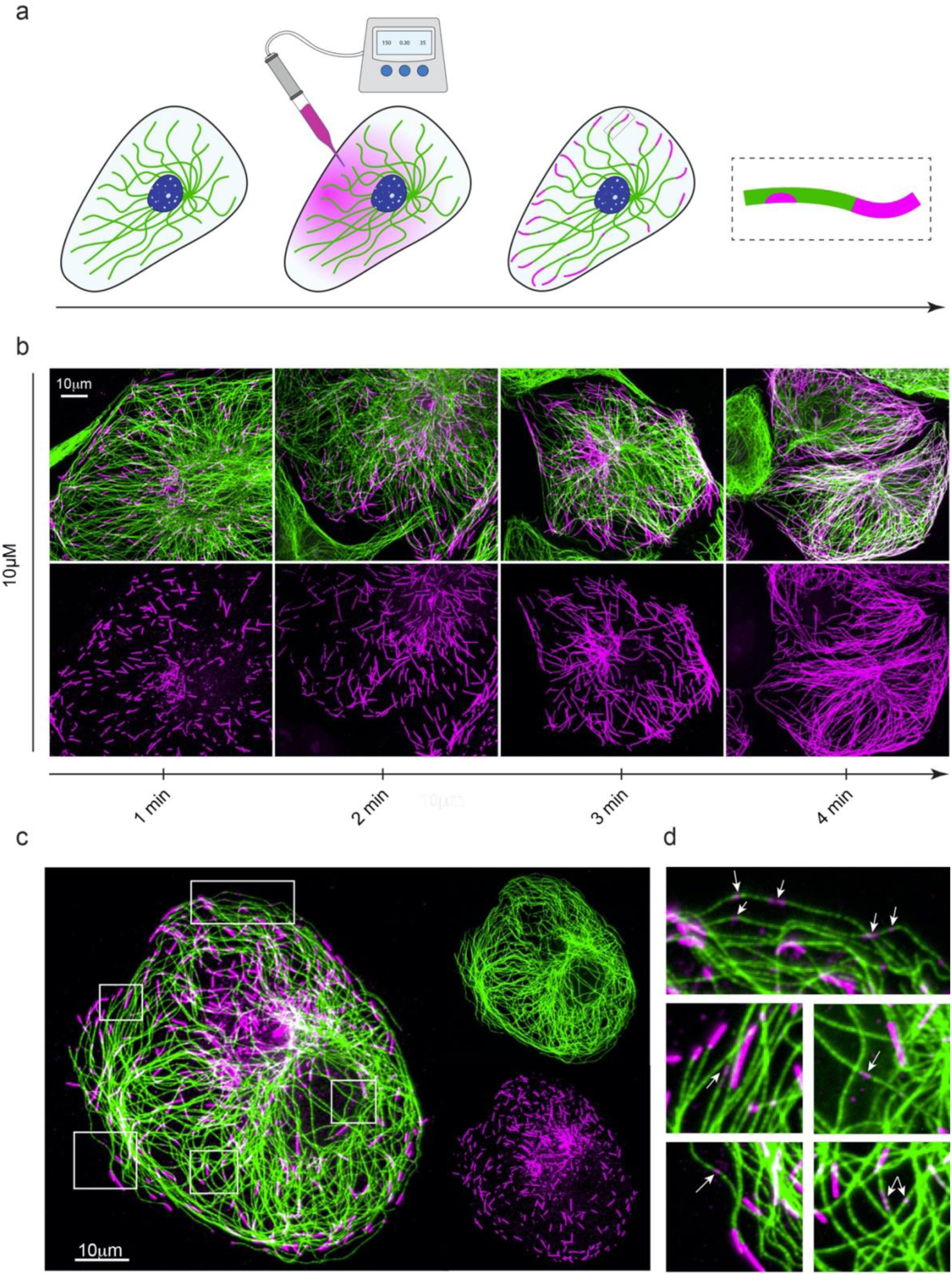
Injected tubulin form patches along microtubules length and polymerize at their growing ends. (**a**) Schematic representation of the microinjection protocol. Purified ATTO-565 labelled tubulin (magenta) is microinjected in PtK2 cells expressing GFP-tubulin (green). Following the microinjection, injected ATTO-565-tubulin (IT) freely diffuse throughout the cell’s cytoplasm. Cells are then pre-permeabilized, fixed and imaged. Right: enlarged region illustrating how IT can for patch along the microtubule’s shaft and assemble at the growing end. (**b**) Self-renewal of microtubule network in PtK2 GFP-tubulin cells injected with 10uM of IT. Z acquisitions were performed and projected on a single image containing the maximal intensity of each pixel. Images show the newly assembled parts of the microtubule network 1,2,3 and 4 minutes post-injection. Scale bar, 10 µm. (**c**) A PTK2 GFP-tubulin cell (ET signal is shown in green) microinjected with 10 µM of IT (IT signal is shown in magenta). Z acquisitions were performed and projected on a single image containing the maximal intensity of each pixel. Scale bar, 10 µm. (**d**) Images correspond to white insets shown in (**c**). Arrows point at IT patches (small magenta spots) that are localized along pre-existing microtubules (green), at distance from microtubules growing ends (long magenta stretches).

### Intensity of newly injected dimers in patches anti-correlates with intensity of pre-existing dimers in the lattice

Following microinjection, microtubules were composed of unlabeled endogenous tubulin dimers (endogenous tubulin), genetically expressed green-fluorescent dimers (Expressed tubulin: **ET**) and microinjected red-fluorescent dimers (Injected tubulin: **IT**) (**Figure 2a**). We reasoned that if the IT dimers replaced the endogenous and the ET dimers, an anticorrelation between the green and the red-fluorescence could be observed. However, if IT dimers were bound to the side of the microtubule lattice, the two intensities should not be related. We thus measured intensity profiles along microtubules displaying red patches along their length. We found that the presence of IT patches along microtubules was associated to a decrease in ET intensities (**Figure 2b**, more examples shown in **Supplementary figure S1a**), supporting the hypothesis of a genuine incorporation in the lattice rather than a side-binding. Moreover, IT intensities measured in IT patches were always lower than the IT intensities measured at microtubule’s growing ends (**Figure 2b** and **Supplementary Figure S1b-c**). This result ruled out the possibility of annealing between the growing ends of two pre-existing microtubules.

**Figure 2:**
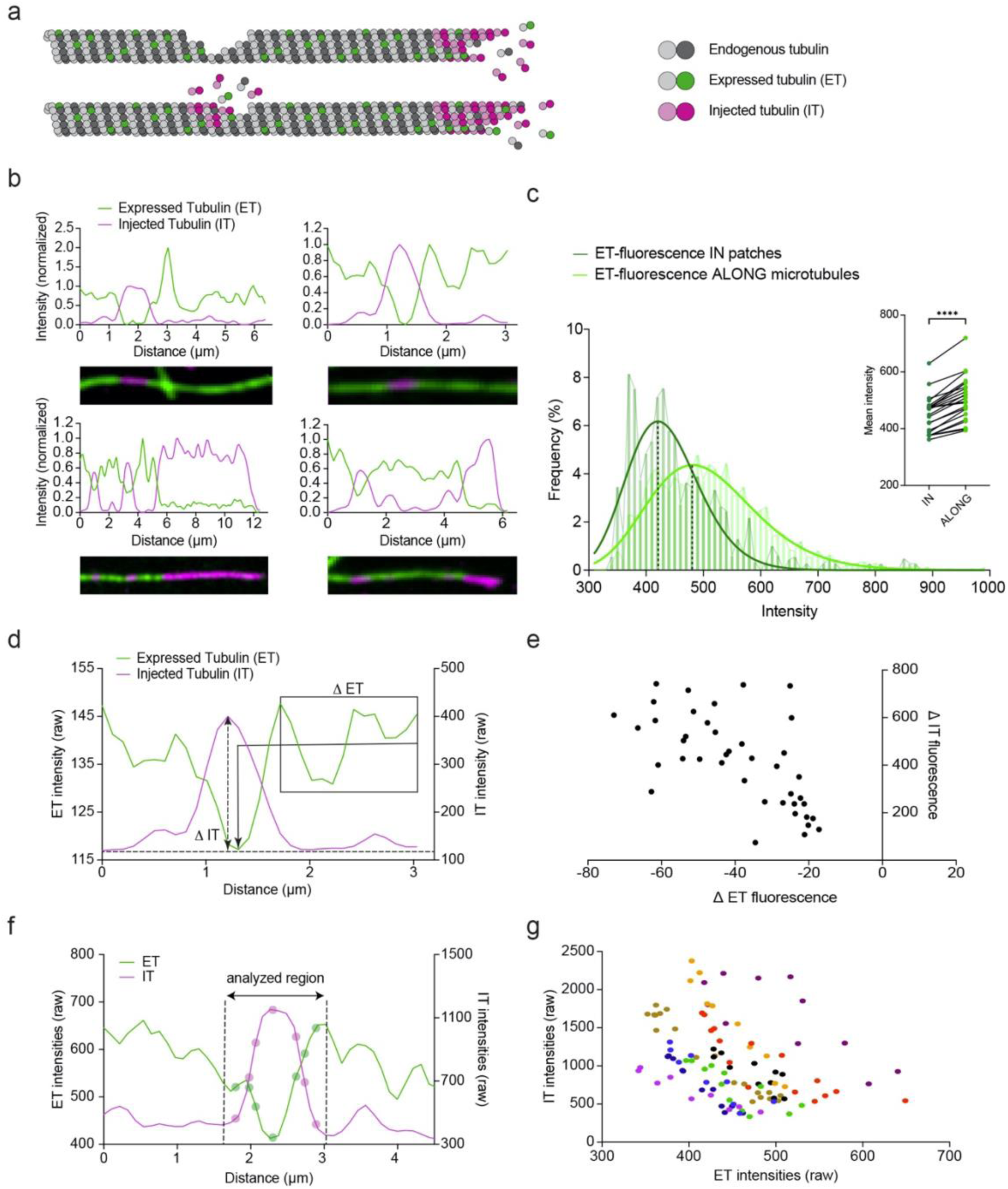
Anti-correlation between injected tubulin patches and fluorescence decrease of pre-existing lattice. (**a**) Schematic representation of the repair process of a damaged microtubule. Following injection of tubulin dimers, the growing end and the damaged region promote the co-assembly of unlabeled endogenous tubulin (grey), GFP-tagged expressed tubulin: ET (green), and ATTO-565-labeled injected tubulin: IT (magenta). (**b**) Fluorescence intensity profiles of microtubules exhibiting a patch of IT. Profiles have been normalized to 1 for the shaft signal of ET and for the peak of IT. (**c**) Histograms of ET-fluorescence intensity frequencies measured in IT patches or in adjacent sections of the microtubule with no detectable IT signal. Black doted lines indicate the means of the two ET fluorescence distributions. Inset represents the differences between the means intensities of ET fluorescence measured in IT patches or in the adjacent section of the microtubule. n=23 from thirteen individual cells. Statistical analysis was performed using two-tailed paired t-test ****p<0.0001. (**d**) Illustration of the measurements of fluorescence intensity variations along the length of a microtubule displaying a patch of IT. The double arrowhead dotted line represents the difference measured between the highest IT-fluorescence intensity and the baseline (DIT). The black square with an arrowhead represents the difference measured between the mean ET-fluorescence intensity along the length of the microtubule and the lowest ET-fluorescence intensity in the patch (DET). (**e**) Graph show the values of DET and DIT measured as described in **d** in PtK2 GFP-tubulin cells microinjected with various concentration of tubulin between 10 and 30 µM. n=23 from thirteen individual cells. (**f**) Illustration of the measurements of fluorescence intensities in a patch of IT. Circles correspond to measurement positions. (**g**) Graph show variations of ET and IT intensities in 9 different linescans measured in cells microinjected with 10, 20 or 30 µM of tubulin (3 linescans per concentration). Distinct colors correspond to distinct microtubules.

To further investigate the decrease in ET intensity in the IT patches, we compared ET intensities measured in IT patches to the ET intensity measured along the entire microtubules. ET intensities in IT patches were consistently lower than ET intensity along the rest of microtubules (**Figure 2c**). The average ET fluorescence measured along IT patches was 11.28 ± 4.84% lower than along microtubules (451.3 ± 64.93 versus 508.7 ± 77.30 mean ± standard deviation) (**Figure 2c**) suggesting that the decrease resulted from a local and specific process rather than from random fluctuations that could be observed elsewhere along the lattice. In addition, the magnitude of the maximal ET decrease (ΔET) measured in the IT patch appeared anti-correlated with the maximal IT increase (ΔIT) (**Figure 2d-e**). Furthermore, point-by-point variations of intensities of the two signals within the patch were also clearly anti-correlated (**Figure 2f-g** and **Supplementary figure S2a-c**). Taken together, our data strongly suggest that injected dimers replaced endogenous and expressed dimers in the lattice, and that observed patches corresponded to genuine incorporations in pre-existing microtubules.

### The incorporation process saturates with time and soluble tubulin concentration

To further investigate the biochemical regulation of the addition of new dimers in a pre-existing lattice, we measured the variations of incorporations intensity as a function of time and concentration. Indeed, a non-specific adsorption of dimers along microtubules should increase linearly with time and tubulin concentration (**Figure 3a-i, 3b**). Conversely, we expect a specific repair mechanism which could be limited to sites where the lattice is damaged, to saturate in time or with the tubulin concentration once damaged sites have been filled up (**Figure 3a-ii, 3b**). We varied IT concentration from 2 to 40 µM and measured the corresponding changes in the IT fluorescence intensity, as well as the length and spatial frequency of incorporations.

**Figure 3:**
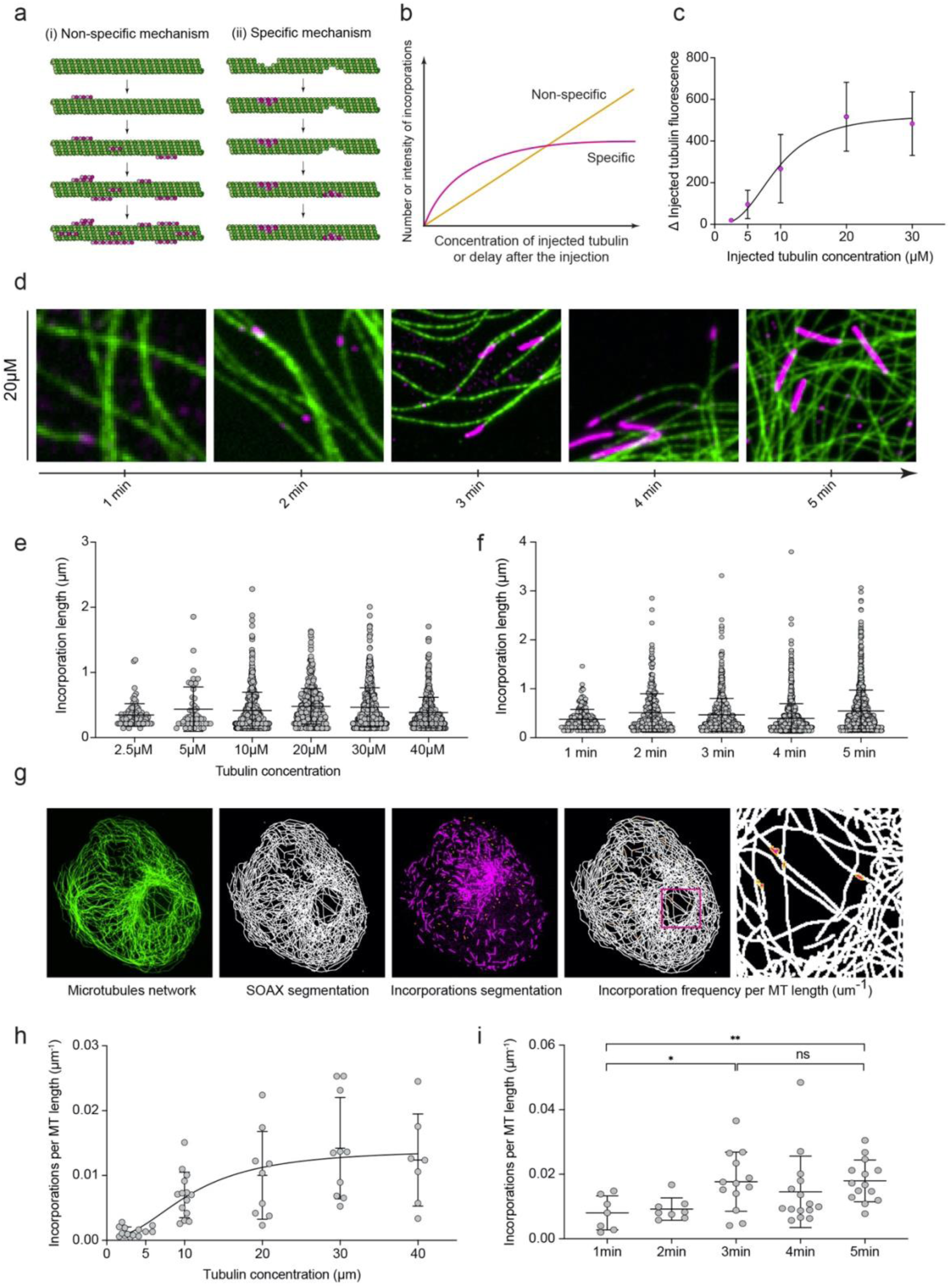
Saturation of incorporations as a function of time and soluble tubulin concentration. (**a**) Scheme depicting two hypotheses for the variations of IT signal (magenta) along pre-existing microtubules (green) over time or with increasing IT concentrations. (i) Non-specific mechanism where IT dimers binds randomly along microtubules. (ii) Self-repair mechanism where IT dimers interacts specifically to damage sites and stop once they are filled up. (**b**) Schematical curves showing the variations of the number or intensity of IT patches with time or increasing IT concentrations according to the two hypotheses illustrated in (**a**). (**c**) Variations of IT intensity in incorporations as a function of IT concentration. Data represents mean ± std.dev., n=9-15 from at least five cells for each condition. (**d**) Representative images of the temporal evolution of IT incorporations (magenta) in pre-existing microtubule network (green) after microinjection. Z acquisitions were performed and projected on a single image containing the maximal intensity of each pixel. (**e**) Incorporation lengths for various IT concentrations. Data represents mean ± std.dev., n=50-471 incorporations measured in each condition in two independent experiments. (**f**) Incorporation lengths depending on the delay after microinjection of 20 µM of tubulin. Data represents mean ± std.dev., n=261-1124 incorporations measured in each condition in two individual experiments. (**g**) Workflow developed to measure the spatial frequency of incorporations in the microtubule network. The microtubule signal is segmented using SOAX (Xu *et al*, 2015) in order to measure the total length of the network. Incorporations are segmented manually. (**h**) Incorporation frequency for various IT concentrations. Data represents mean ± std.dev., n = 5-14 analyzed cells in each condition in two independent experiments. (**i**) Incorporation frequency depending on the delay after microinjection of 20 µM of tubulin. Data represents mean ± std.dev., n=7-15 cells analyzed for each condition in two independent experiments. Statistical analysis was performed using two-tailed unpaired t-test. *p=0.0202, **p=0.0024.

The proportion of IT in microtubule’s growing ends increased progressively with the IT concentration (**Supplementary Figure S3.a-d**), confirming that variations of cytoplasmic concentration corresponded to our manipulation of injected concentrations. Interestingly, IT intensity in incorporations increased and then reached a plateau beyond 20 µM of IT (**Figure 3c**). In parallel, the decrease of ET intensity in incorporations was again anti-correlated to the IT signal (**Supplementary Figure S3e**).

Interestingly, the length of incorporations did not vary with IT concentration (**Figure 3e**) nor with the delay post-injection (**Figure 3f**) in the ranges we tested. These results suggested that damaged sites were filled with injected dimers in less than a minute and did not elongate in the next few minutes.

We then designed a workflow to measure accurately the spatial frequency of incorporations in the network (**Figure 3g**). This required not only the identification of repair sites, but also an evaluation of the entire length of the microtubule network. Using the SOAX software (Xu *et al*, 2015), we could accurately segment the microtubule network. Incorporations were then segmented manually by selecting the patches whose IT intensities were at least twice higher than the background signal along microtubules. The spatial frequency of incorporations per cell thus corresponded to the total number of incorporations divided by the length of the entire microtubule network. By varying IT concentration, we found that the spatial frequency of incorporations saturated beyond 20 µM (**Figure 3h**). Interestingly, this saturating concentration of 20 µM also corresponded to the value beyond which the intensity of individual incorporations reached a maximum (**Figure 3c**). At this concentration, the frequency of incorporations increased during the first 3 min after the injection and then saturated (**Figure 3i**). Altogether, these results point at a saturation of the incorporation process, supporting the hypothesis of specific repair mechanism that is limited to a defined number of sites rather than a non-specific mechanism all along the length of microtubules. We could thus evaluate the frequency of microtubule repair in saturating conditions (20 µM of IT, 3 minutes post-injection) to be around 15 sites per mm of microtubules in PtK2 cells, i.e., a repair site every 70 microns (**Figure 3h-i**).

### Repair sites are not evenly distributed in the cell

Noteworthy, we observed that incorporations were longer in the vicinity of the injection site than in the cell periphery (**Supplementary Figure S4a-d**). This showed that the injection flow can induce some damages along microtubules, as shown previously *in vitro* (Portran *et al*, 2017; Schaedel *et al*, 2015). Fortunately, these experiments also showed that the injection flow did not propagate beyond 20-30 µm from the injection site (**Supplementary movie S1**), and that repair sites more distant than this critical length were not affected by this flow (**Supplementary Figure S4b**). This showed that despite the damaging effect of the injection flow, the localization of repair sites and the variations of their subcellular density could be investigated in living cells. At first glance, it seemed that they were not evenly distributed all over the cell (**Supplementary Figure S4b**), suggesting that the repair process did not occur randomly on all microtubules.

To describe quantitatively the spatial distribution of incorporations in the microtubule network, we normalized cell shapes and internal architectures using micropatterns. Crossbow-shaped micropatterns direct the assembly of a lamella (transverse arcs and radial fibers) along the adhesive and curved edge (**Figure 4a**, upper part of the image), and stress fibers along non-adhesive and straight regions (**Figure 4a**, upper part of the image) (Théry *et al*, 2006). These actin structures are reproducible over many cells and do not fluctuate much in a period of 3 minutes (**Figure 4b-c, Supplementary movie S2**). The microtubule network was denser at the cell center and along the adhesive and curved edges, and less dense close to stress fibers (**Figure 4d-e**). Interestingly, although lateral fluctuations of microtubules could be observed throughout the cell cytoplasm, they were slightly more pronounced in the vicinity of stress fibers (**Figure 4f, Supplementary movie S3**).

**Figure 4:**
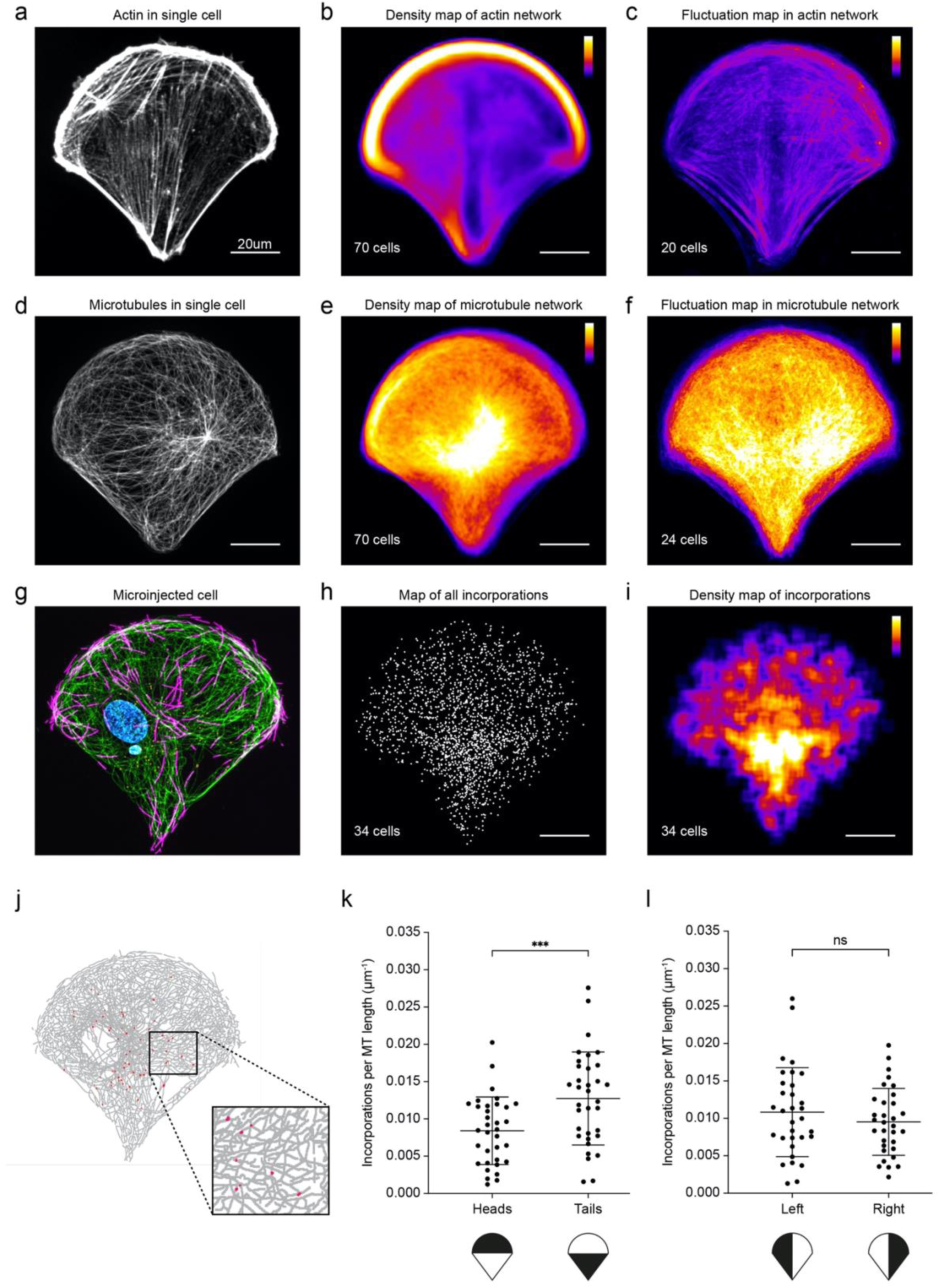
Distribution of repair sites in micropatterned PtK2 cells. (**a**) Representative image of the actin network in PtK2 GFP-tubulin cell plated on a crossbow-shape fibronectin-coated micropattern. Z acquisitions were performed and projected on a single image containing the maximal intensity of each pixel. Scale bar, 10 µm. (**b**) Averaged fluorescence signal of F-actin of 70 images. Intensities are color coded with the “fire” look-up table. Scale bar, 10 µm. (**c**) Averaged fluctuations of actin signal reporting the subcellular regions where actin bundles move. Actin signal was measured every 4 seconds for 3 minutes. The standard deviation of the signal was measured for each individual cells and averaged over n=20 cells. Intensities were color coded with the “fire” look-up table. Scale bar, 10 µm. (**d**) Representative image of the microtubule network in PtK2 GFP-tubulin cell. Z acquisitions were performed and projected on a single image containing the maximal intensity of each pixel. Scale bar, 10 µm. (**e**). Averaged fluorescence signal of microtubules (GFP signal) of 70 images. Intensities are color coded with the “fire” look-up table. Scale bar, 10 µm. (**f**) Averaged fluctuations of tubulin signal reporting regions where microtubules move. Tubulin signal was measured every 10 seconds for 3 minutes. The standard deviation of the signal was measured for each individual cells and averaged over n=20 cells. Intensities were color coded with the “fire” look-up table. Scale bar, 10 µm. (**g**) Representative image of a PtK2 GFP-tubulin (green) cell microinjected with ATTO-565 tubulin (magenta). Z acquisitions were performed and projected on a single image containing the maximal intensity of each pixel. Scale bar, 10 µm. (**h**) Overlay of the positions of incorporations measured in 34 cells from six independent experiments. Scale bar, 10 µm. (**i**) Density map of incorporations measured in **h**. The distribution of positions was spatially averaged using a 40-pixel-wide sliding square and color coded with the “fire” look-up table. Scale bar, 10 µm. (**j**) Representative image of a SOAX segmented microtubule network (grey) containing IT incorporations (red) in a PtK2 GFP-tubulin cell micropatterned on a crossbow shape. (**k**) Spatial frequency of incorporations measured in the top and bottom part of PtK2 cells micropatterned on crossbow shapes. Data represents mean ± std.dev., n=34 cells from six individual experiments. Statistical analysis was performed using two-tailed paired t-test ***p=0.0005. (**l**) same as (k) but measurements were performed in the left and right parts of the cells.

We microinjected red-fluorescent tubulin dimers in micropatterned cells to assess the localization of repair sites (**Figure 4g**). By reporting all the positions of incorporations detected over 34 cells (**Figure 4h**), we could generate a density map of their localization (**Figure 4i**). Strikingly, the density map of incorporations (**Figure 4i**) did not correspond exactly to the density map of microtubules (**Figure 4e**). Incorporations were rarely observed along adhesive and curved edge where the network is dense. They were more frequent close to cell center and seemed biased toward the regions adjoining the stress fibers. However, microtubule network density was quite high in those regions and could have biased the evaluation of local repair density. In addition, average densities of repair sites could be affected by cell-to-cell variations in the microtubule network architecture. Therefore, it required to measure local spatial frequencies in individual cells. To that end, we segmented microtubules and repair sites in individual cells (**Figure 4j**). We compared repair densities in regions facing distinct actin network architectures, and found that they were more frequent toward the stress fibers (12.7 ± 6.2 per mm of MT) than toward the lamella (8.4 ± 4.5 per mm of MT) (**Fig. 4k**). As a control, we compared the left and the right part of the cell, symmetric in terms of cell adhesion and actin architecture, and found no difference (10.8 ± 5.9 and 9.5 ± 4.5 incorporation per mm of MT) (**Fig 4l**). Interestingly, microtubule fluctuations were more intense in those regions, suggesting that repair could be induced by local bending forces. Altogether, these results further confirmed the specificity of the repair process by revealing that it was not evenly distributed over the entire network. We concluded that incorporations were more frequent in regions adjoining stress fibers, where cortical actin was less dense and where microtubules displayed higher lateral fluctuations.

## Discussion

Our study provides the first experimental evidence that microtubules walls undergo self-repair in living cells. The negative correlation measured between the IT and ET intensities demonstrates that newly injected dimers are embedded in the microtubule lattice. Furthermore, the lower IT intensities in incorporations than microtubule tips discard the possibility of annealing between two microtubule growing ends. Finally, the saturation of IT intensities in incorporations, the saturation of the incorporation frequency as a function of time and concentration, and their preferential location in defined subcellular regions attest for the non-random binding of IT dimers in the pre-existing microtubule network. Altogether, our results attest the specificity of tubulin incorporations in limited number of sites in the microtubule lattice that are not randomly localized along microtubules. The preferential location of these sites in regions where microtubules displayed large lateral fluctuations and the saturation of the polymerization process strongly suggest that these incorporations are genuine repair sites where damaged lattice is healed by the addition of free dimers.

Interestingly, the ratio between the fluorescence intensity in incorporations and growing ends (**Supplementary Figure S1c**) suggested that these incorporations contained between 4 and 8 repaired protofilaments. Smaller damages, in which only few protofilaments were repaired, might have been too dim to pass the fluorescence threshold we imposed (twice the background) to be counter as incorporation sites. On the contrary, it is possible that larger damages involving a higher number of protofilaments could not be repaired and rather triggered microtubule breakage and disassembly. In any case, it is interesting to note that about half of the section of the microtubule could be destroyed and repaired. This value is much larger than the low fluorescence intensity of incorporations observed along non stabilized microtubules *in vitro* (1 to 10% of lattice dimers) (Schaedel *et al*, 2019). It would be interesting to know more about the contribution of specific microtubule-associated proteins to ensure the survival of such large damages and to their expression specific cell types in which microtubules are more prone to be injured.

The value of one repair sites every 70 µm of microtubules (**Figure 3h-i**) is likely an underestimation of the actual spatial frequency of the repair sites. First, we considered only large damage sites in which IT fluorescence was twice higher than the background, so we discarded all the small repair involving few dimers. Second, the polymerization of IT at microtubules growing ends prevented us from looking at the repair process over duration longer than five minutes. Considering that microtubule’s lifetime varies between 10 and 30 minutes (data not shown), it’s possible that the total number of incorporations during the entire lifetime of the microtubule is higher than our measurement. Finally, some repair sites may also have been protected by microtubule-associated proteins such as CLASP or SSNA1 without involving the incorporation of tubulin dimers (Aher et al, 2020; Lawrence et al, 2021). Overall, the quantifications provided in this study report the spatial frequency of large damages in the lattice that have been repaired in a limited time window and involved dimer replacement. They likely largely underestimated the actual dynamics of the microtubule lattice.

Finally, we observed that incorporations were more frequent in regions adjoining stress fibers than in the lamella. It is counter-intuitive that the microtubules that were bent by the retrograde flow of the actin network in the lamella did not display more damages and repairs. It is possible that the characteristic timescale of actin retrograde flow (several dozen of minutes) was too long to generate damages and repair in the time window we analyzed (5minutes). At this time scale, the actin network barely changed in the lamella (**Figure 4c**). The mechanism responsible for the incorporations we observed might operate at shorter time scales. Our investigation of microtubule network fluctuations, by monitoring microtubule shapes every 10s during 5 minutes, revealed an interesting correlation with the spatial distribution of repair sites (**Figure 4f** and **4i**). It is not yet known whether these fluctuations are at the origin of the microtubule damage or whether both phenomena are independent manifestations of the same originating mechanism. Indeed, molecular motors can both deform (Jolly *et al*, 2010; Randall *et al*, 2017) and destroy microtubules (Triclin *et al*, 2021). In a nonexclusive mechanism, bending forces could also directly induce microtubule destruction and repair (Schaedel *et al*, 2015). Whether and how the regulation of microtubule damage and repair has any impact on microtubule stability and cell polarity (Aumeier *et al*, 2016; Andreu-Carbó *et al*, 2021) is an interesting hypothesis that deserve further investigation.

Altogether, our results demonstrated that microtubules undergo frequent self-repair in living cells. They also revealed the existence of sub-cellular regulation mechanism that might contribute to determine specific populations of microtubules. The physiological impact of this destruction and repair mechanism remains to be determined.

## Materials and methods

### Cell culture

Male rat kangaroo kidney epithelial cells (PtK2) stably expressing GFP-Tubulin obtained from Franck Perez lab were grown at 37°C and 5% CO_2_ in DMEM/F12 (31331028, Gibco) supplemented with 10% fetal bovine serum (10270106, Life Technologies) and 1% antibiotic-antimycotic solution (15240062, Gibco). The day before experiments, cells were detached using TrypLE (12605036, GIBCO) and plated in glass bottom dish (627860, Dutscher) or in micropatterned coverslips glued to home-made cut out plastic dish.

### Tubulin purification and labelling

Fluorescent tubulin (ATTO-565-labelled tubulin) were prepared as previously described (Shelanski, 1973; Hyman *et al*, 1991; Vantard *et al*, 1994). Briefly, tubulin was purified from fresh bovine brain by three cycles of temperature-dependent assembly and disassembly in Brinkley buffer 80 (BRB80 buffer: 80mM PIPES pH 6.8, 1mM EGTA and 1mM MgCl2 plus 1mM GTP). MAP-free neurotubulin was purified by cation-exchange chromatography (EMDSO, 650M, Merck) in 50mM PIPES, pH 6.8, supplemented with 1mM MgCl2 and 1mM EGTA. Purified tubulin was obtained after a cycle of polymerization and depolymerization. Finally, tubulin was labelled with ATTO-565 fluorochromes, resuspended in microinjection buffer (50 mM potassium glutamate, 1 mM MgCl2, pH 6.8), aliquoted in 500µl plastic tubes, flash frozen in liquid nitrogen and store at -80°C.

### Microinjection

Glass microneedles were home-made pulled from clark borosilicate thin wall capillary (30-0050 Harvard Apparatus) using a horizontal pipet puller (PN-3, Narishige). Purified labelled tubulin was thaw and diluted in injection buffer (50 mM potassium glutamate, 1 mM MgCl2, pH 6.8) on ice to reach final concentration. Microneedles were loaded with 5µL of tubulin solution and connected to a FemtoJet 4i (Eppendorf). Microneedles were manually controlled with an InjectMan 4 micromanipulator (Eppendorf). The compensation pressure set on the FemtoJet was 35hPa for the whole injection experiment to avoid damages on cells. Microinjection of PTK2 cells were performed in an inverted microscope (Nikon Ti2 Eclipse) equipped with a Prime BSI Express CMOS camera (Photometrics) and using a Nikon CFI Plan Fluor 40x/0.75 NA dry objective. Cell medium was maintained at 37°C and 5% CO2 during the whole experiment using a H-301 heating chamber (Okolab). Micro-Manager 1.4.21 software was used for images acquisition.

### Cells fixation and labeling

Before fixation, cells were permeabilized with BRB80 1X supplemented with 0.25% Triton-X100 (T8787, Sigma) for 30 sec, and then fixed in cytoskeleton buffer (MES 10mM, KCl 138mM, MgCl 3mM, EGTA 2mM) supplemented with 10% sucrose, 0.5% Triton-X100 and 0.5% glutaraldehyde (G5882, Sigma). Aldehyde functions were then reduced using 1mg/ml of NaBH_4_ for 10 minutes at room temperature. Cells were washed 3 times with PBS-Tween 20 (1379, Sigma) 0.1% and incubated with phalloidin for 30 min and DAPI (D9542, Sigma) for 5 minutes. Coverslips were finally washed 3 times in PBS-Tween 20 0.1% and mounted in Mowiol 4-88 (81381, Sigma).

### Imaging

Immunofluorescence images of GFP microtubules and injected ATTO-565-tubulin in cells were acquired using a confocal spinning disk microscope (Nikon Ti Eclipse equipped with a spinning scanning unit CSU-X1 Yokogawa) and a R3 retiga camera (QImaging). Images were acquired using a Nikon Plan Apo VC 60x/1.40 NA oil objective using a 1.5x additional integrated magnification lens. Each wavelength was acquired separately with a 200nm Z-step width. Metamorph software was used for images acquisition.

### Micropatterning

Clean glass coverslips were activated with an oxygen plasma treatment (PE50 - PlasmaEtch) for 30s at 30W and incubated with 0.1 mg/ml of poly(L-Lysine)-poly(ethylene-glycol) (PLL-PEG JenKem Technology) in 10 mM Hepes, pH 7.4, at room temperature for 1 h. Coverslips were then dried following dewetting in the presence of 1ml of ultra-pure H_2_O. PLL-PEG coated coverslips were placed in contact with an optical mask containing transparent micropatterns (Toppan Photomasks, Inc.) using a home-made vacuum chamber and exposed for 4 min to deep UV light (UVO Cleaner 42; Jetlight Company). Micropatterned slides were finally incubated for 20 min with a solution of 10 µg/ml bovine fibronectin solution (F1141, Sigma) and 10 µg/ml of Collagen I Rat Protein, Tail (A1048301, GIBCO) in NaHCO_3_ 10mM. Before plating cells, patterned coverslips were washed one time with NaHCO_3_ followed by three washes with ultra-pure H_2_O.

### Image and statistical analysis

Acquired images were analyzed using ImageJ. In the red channel of acquired images (corresponding to the injected tubulin), 3 squares at different cell locations were chosen to measure the mean fluorescent intensity of the background. Red signal not corresponding to growing tips, and above 2 time the background signal on MT were manually selected as incorporations. The total MT length was obtained using the SOAX software (Xu *et al*, 2015). Finally, the incorporation frequency was measured by dividing the number of incorporations on the total length of the MT network. For the density maps, actin and microtubules images were aligned using the StackReg plugin in imageJ, projected on a single image containing the average intensity of each pixel and intensities were color coded with the “fire” look-up table. For the fluctuation maps, the background of the actin and microtubule images was subtracted, and they were equalized for the dynamic ranges of grey values. The standard deviations of each film were then stacked, aligned using StackReg plugin in imageJ, and projected on a single image containing the average intensity of each pixel. The intensities were color coded with the ‘fire” look-up table. Statistical analysis was performed using GraphPad Prism software (version 9.0), a p-value above 0.05 was consider as significant.

## Acknowledgements

This work was supported by the European Research Council (Consolidator Grant 771599 (ICEBERG) to MT and Advanced Grant 741773 (AAA) to LB), by the Bettencourt-Schueller foundation, the Emergence program of the Ville de Paris and the Schlumberger foundation for education and research. This project was also supported by the MuLife imaging facility, which is funded by GRAL, a program from the Chemistry Biology Health Graduate School of University Grenoble Alpes (ANR-17-EURE-0003).

---

**Figure S1:**
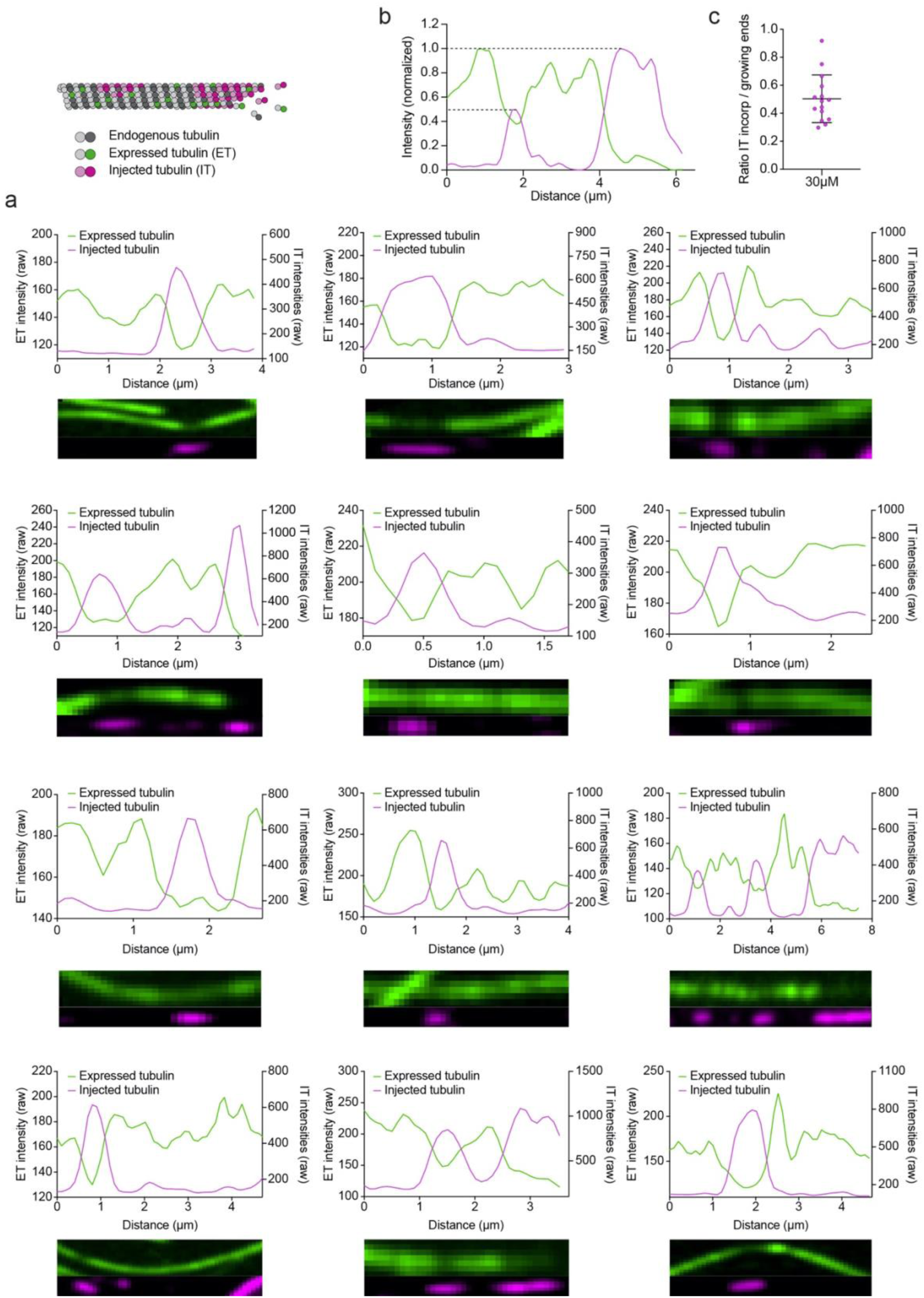
Examples of anti-correlation between injected tubulin patches and fluorescence decrease of pre-existing lattice. (**a**) Fluorescence intensity profiles of microtubules exhibiting a patch of IT. Corresponding fluorescence images of expressed tubulin (ET) and injected tubulin (IT) are shown in green and magenta, respectively. (**b**) Representative fluorescence intensity profiles of microtubules exhibiting a patch of IT. Profiles have been normalized to 1 for the shaft signal of ET and for the peak of IT. (**c**) IT-fluorescence intensity ratios between the maximum intensity measured in the patches and the maximum intensity measured at the growing ends in PtK2 GFP-tubulin cells microinjected with 30 µM of IT. Data represents mean ± std.dev, n=15 intensity profiles from six different cells.

**Figure S2:**
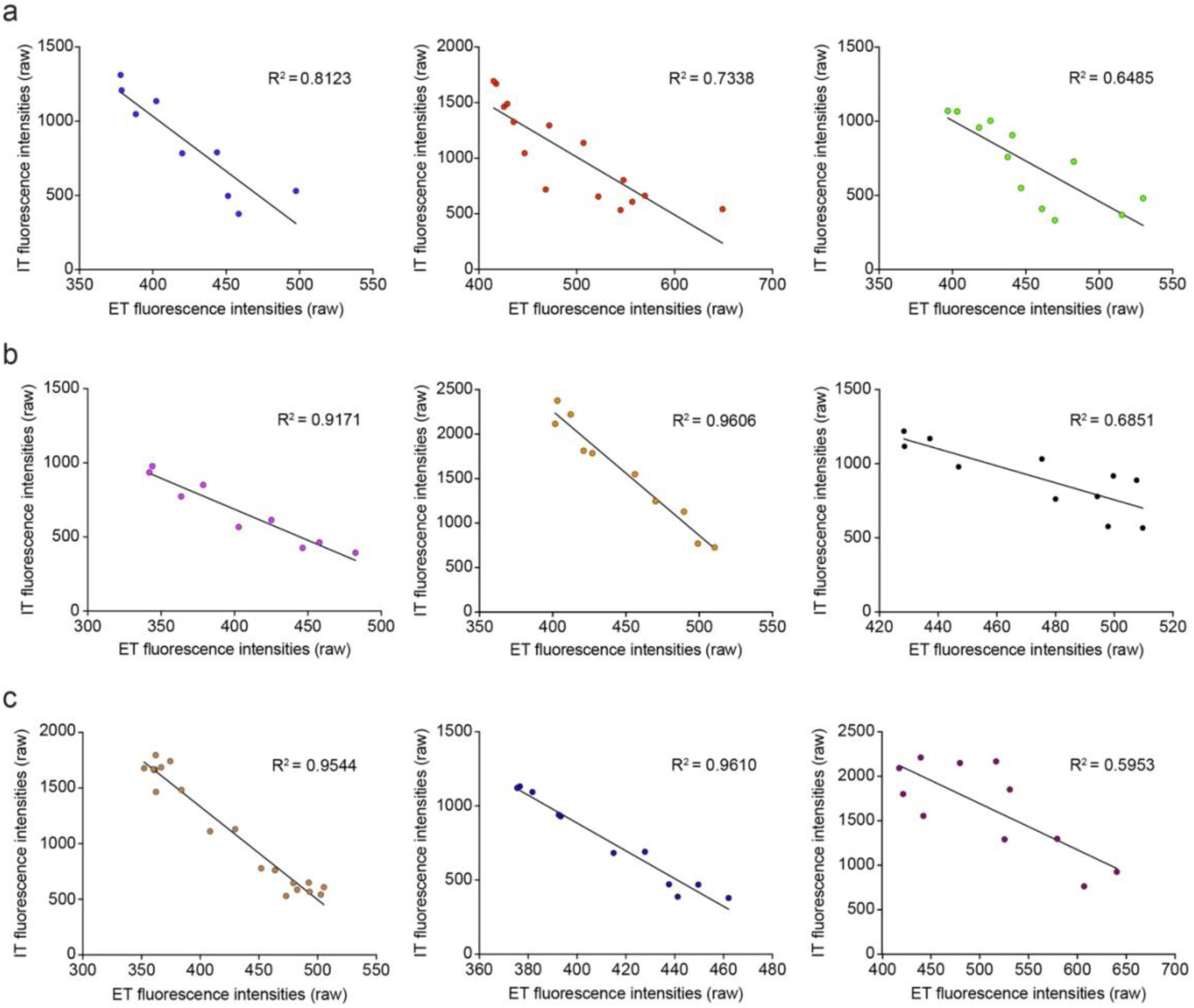
Anti-correlation between injected tubulin intensity and endogenous tubulin intensity in patch of injected tubulin. (**a**) Correlations between injected tubulin intensities and expressed tubulin intensities measured in three different linescans in cells microinjected with 10 µM of tubulin. R squared were measured using linear regression. (**b**) Correlations between injected tubulin intensities and expressed tubulin intensities measured in three different linescans in cells microinjected with 20 µM of tubulin. R squared were measured using linear regression. (**c**) Correlations between injected tubulin intensities and expressed tubulin intensities measured in three different linescans in cells microinjected with 30 µM of tubulin. R squared were measured using linear regression.

**Figure S3:**
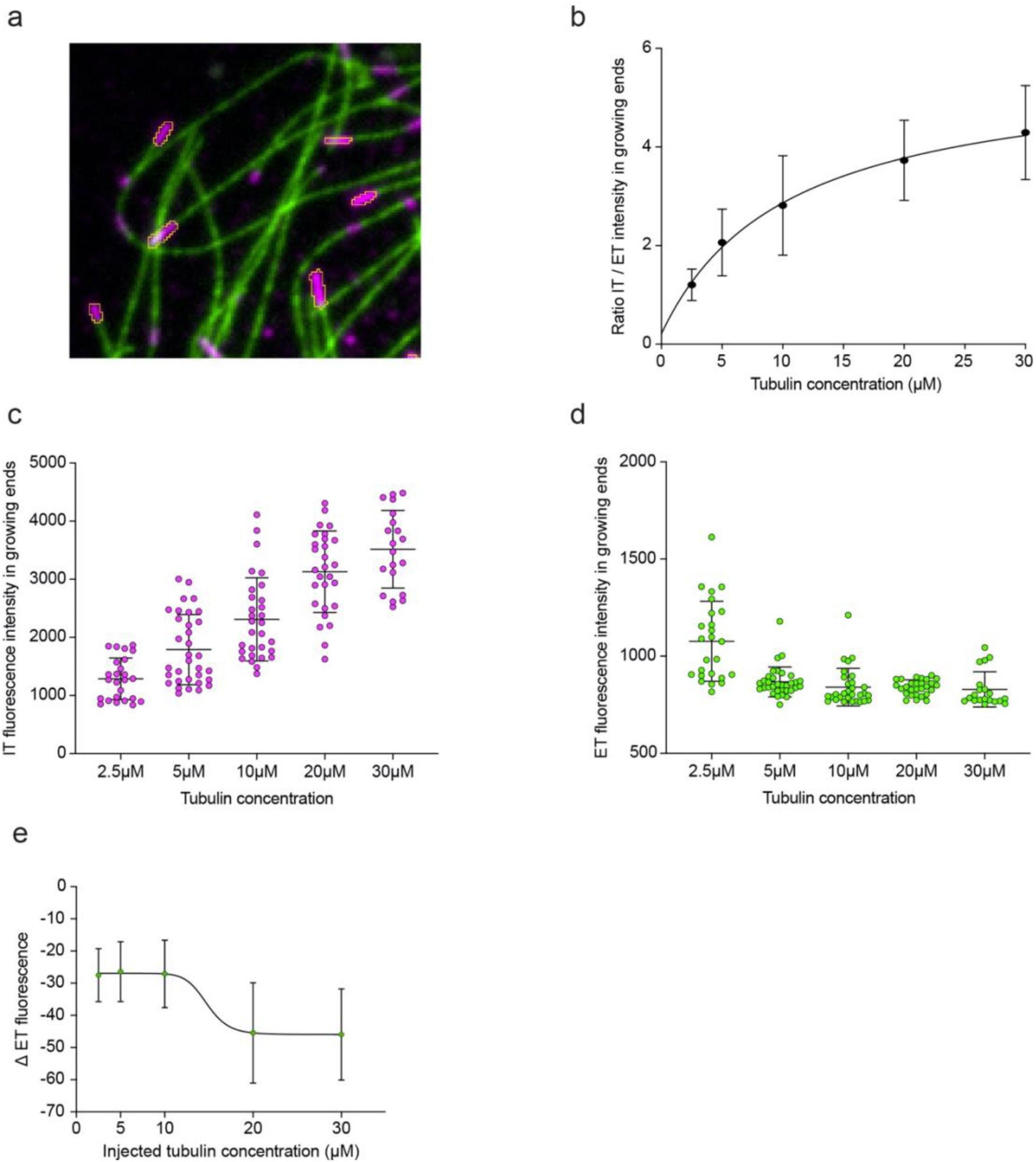
Amount of injected tubulin at growing ends and damage sites is concentration dependent. (**a**) Representative image of microtubule growing ends selection. Z acquisitions were performed and projected on a single image containing the sum intensity of each pixel. IT and ET fluorescence intensities are measured in the selected regions. (**b**) Fluorescence intensity ratios between IT and ET measured at microtubule growing ends. (**c**) IT fluorescence intensity measured at microtubule growing ends. (**d**) ET fluorescence intensity measured at microtubule growing ends. In **b, c** and **d**, each point corresponds to the mean of ± 30 microtubule growing ends in a cell. Data represents mean ± std.dev, n=25-33 cells for each condition from at least four individual experiments. (**e**) Intensity variations of the expressed tubulin in incorporations as a function of the injected concentration. Data represents mean ± std.dev., n=9-15 from at least five cells for each condition.

**Figure S4:**
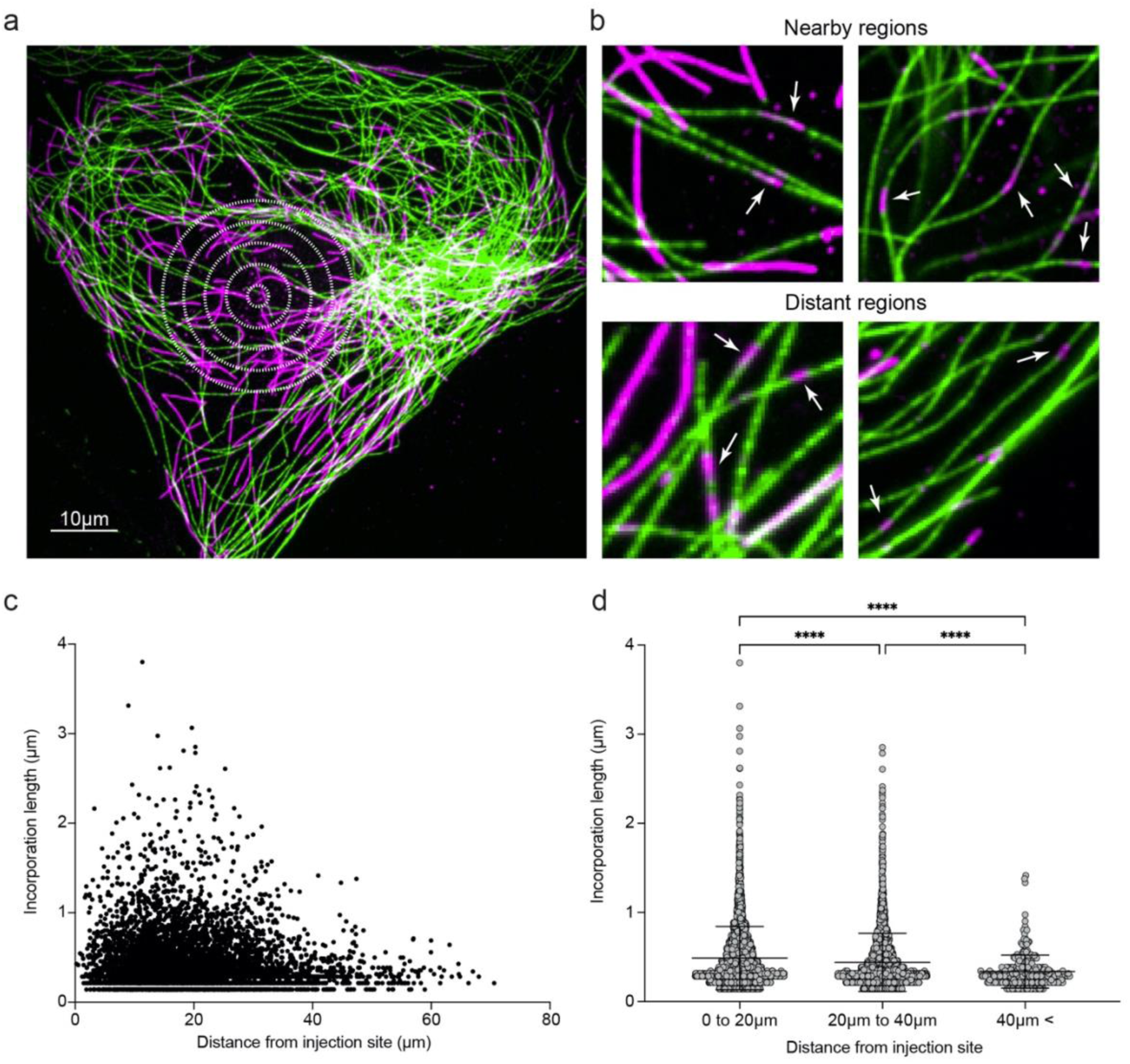
Distribution of incorporation length depending on the distance to the injection site. (**a**) Representative image of PtK2 GFP-tubulin cell microinjected with 10 µM of tubulin. White concentric circles are representing the injection site. Scale bar, 10 µm. (**b**) Enlarged regions of nearby and distant regions from two different cells. White arrows are showing the presence of magenta IT incorporations in the pre-existing microtubule network. (**c**) Dot plot of incorporation lengths as a function of the distance from the injection site. (**d**) Incorporation lengths measured as a function of the distance from the injection site grouped by 20µm size ranges. In **c** and **d**, n=5703 incorporations measured from 111 cells used in the whole study.

**Supplementary movie 1: Microinjection’s effect on the microtubule network**.

Visualization of the effect of injection flow (buffer without tubulin in order to monitor the pre-existing network properly) on the microtubule network of a PtK2 cell expressing GFP-tubulin (shown in white). Images were recorded every 0.1 second for 8 seconds with a 60x oil objective on an inverted microscope.

**Supplementary movie 2: Representative actin dynamic for 3 minutes in a micropatterned crossbow cell**.

Monitoring of the evolution of the actin network at short time scales (3 minutes). Actin filaments were labelled with SiR-Actin (show in grey) in a PtK2 cell (expressing GFP-tubulin, not shown) plated on a crossbow-shaped fibronectin-coated micropattern. Images were recorded every 4 seconds for 180 seconds with a 60x oil objective on a spinning disk confocal microscope.

**Supplementary movie 3: Representative microtubule dynamic for 3 minutes in a micropatterned crossbow cell**.

Monitoring of the evolution of the actin network at short time scales (3 minutes). PtK2 cell expressing GFP-tubulin (shown in white) were plated on a crossbow-shape fibronectin-coated micropattern. Images were recorded every 10 seconds for 180 seconds with a 60x oil objective on a spinning disk confocal microscope.

